# Rapid antimicrobial susceptibility test using spatiotemporal analysis of laser speckle dynamics of bacterial colonies

**DOI:** 10.1101/853168

**Authors:** SeungYun Han, Hojun No, YoonSeok Baek, Huijun Park, KyeoReh Lee, Seungbum Yang, YongKeun Park

## Abstract

Antimicrobial susceptibility testing (AST) is crucial for providing appropriate choices and doses of antibiotics to patients. However, standard ASTs require a time-consuming incubation of about 16-20 h for visual accumulation of bacteria, limiting the use of AST for an early prescription. In this study, we propose a rapid AST based on laser speckle formation (LSF) that enables rapid detection of bacterial growth, with the same sample preparation protocol as in solid-based ASTs. The proposed method exploits the phenomenon that well-grown bacterial colonies serve as optical diffusers, which convert a plane-wave laser beam into speckles. The generation of speckle patterns indicates bacterial growth at given antibiotic concentrations. Speckle formation is evaluated by calculating the spatial autocorrelation of speckle images, and bacterial growth is determined by tracking the autocorrelation value over time. We demonstrated the performance of the proposed method for several combinations of bacterial species and antibiotics to achieve the AST in 2-4.5 hours. Furthermore, we also demonstrated the sensitivity of the technique for low bacterial density. The proposed method can be a powerful tool for rapid, simple, and low-cost AST.

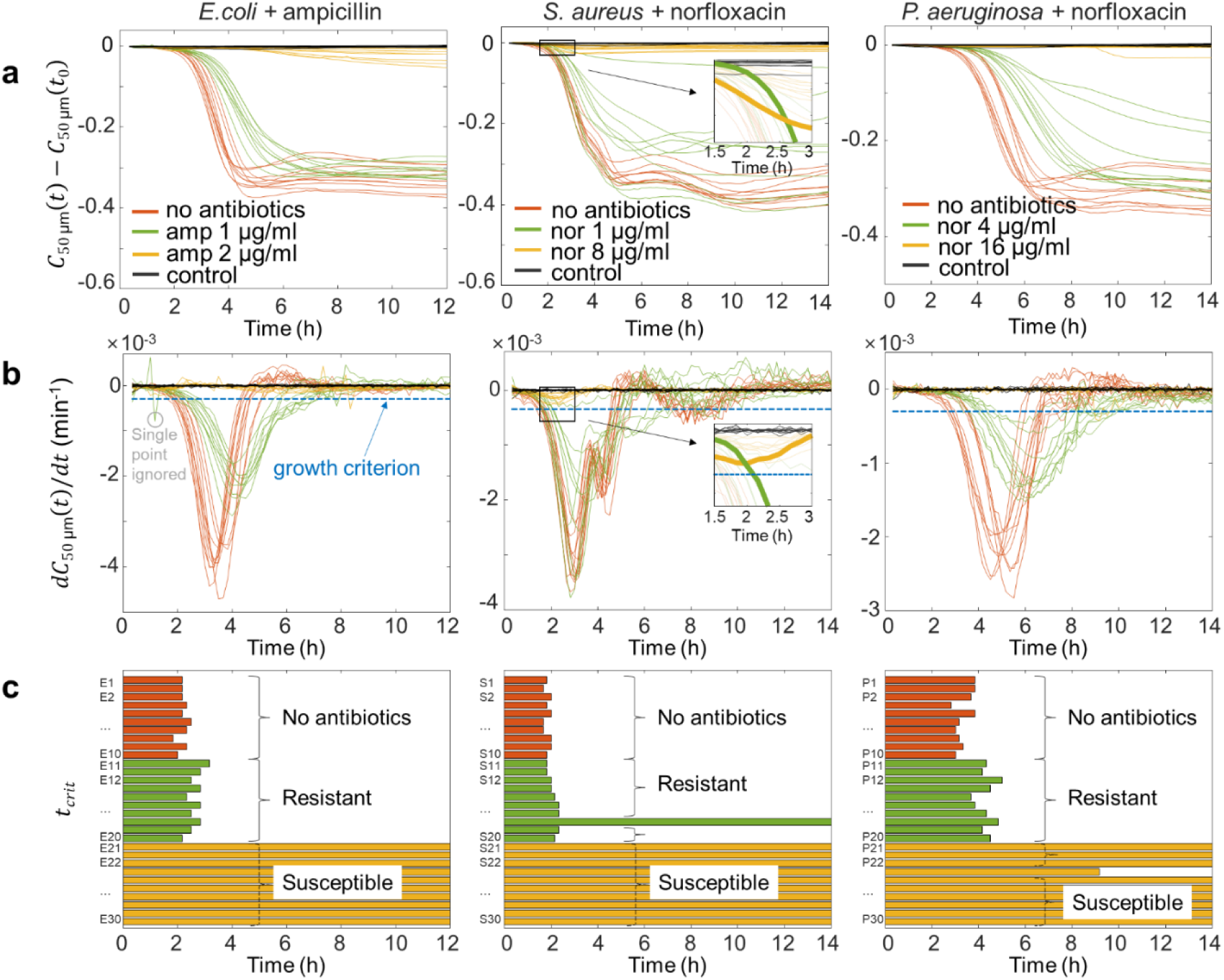

## INTRODUCTION

The misuse and overuse of antibiotics have increased the incidence of antimicrobial resistance, posing a great threat to global health care by reducing the efficacy of antibiotics for treating pathogen-based diseases^1^. Antimicrobial susceptibility testing (AST) is thus necessary to prevent the improper use of antibiotics. It is a routine clinical procedure to determine the susceptibility of pathogens to given antibiotics and their doses. AST provides information on the efficacy of certain antimicrobial agents to a pathogen, thus helping physicians to choose an ideal treatment for a patient. However, utilizing the desired information using standard AST methods requires a lead time longer than one day^2-4^, mainly due to the culturing process of bacteria. Solid-based ASTs, such as agar dilution, disk diffusion, and E-test, use visual detection of bacterial colony growth, which requires 16-24 hours^5^. Liquid-based tests such as broth microdilution or automated instruments to measure the optical density (OD) of a bacterial solution evaluate growth at a given antibiotic condition^6-11^. These tests require an incubation time of 6-8 hours for the production of a high-density bacterial solution because of their low sensitivity for OD measurement^12^. This long lead time hinders the availability of AST results when physicians choose the initial prescription for patients. Thus, physicians are often required to offer initial antibiotics based on their empirical guess^13^. Such a process includes the chance of prescribing improper antibiotics and their doses, resulting in the emergence of antibiotic-resistant bacteria. Hence, a more rapid AST that can offer an early guide to physicians is of urgent need to prevent the crisis of antibiotic resistance.

To achieve rapid AST, numerous diagnostic techniques have been developed for decades. Many approaches exploit single-cell optical imaging to monitor cellular growth at different antibiotic conditions. These techniques use time-lapse images of a single bacterial cell immobilised in various ways such as agarose gel^12^, microfluidic chip^14,15^, confined microchannels^16^, or nanolitre arrays^4^. However, these methods rely on high-resolution imaging and time-lapse investigation of multiple locations, which makes them expensive and complicated for high-throughput performance. Non-optical approaches utilise cell mass changes using magnetic bead rotation^17^ or cantilevers^18^, fluorescent signals from the metabolic activity of bacteria^19^, or rapid-growth agar^20^. These techniques also require complex readout systems, and their compatibility with standard methods should be demonstrated for point-of-care use. Biochemical ASTs have also been suggested using Raman spectroscopic biomarkers^21^, DNA probes^22,23^, and RNA markers^24^. Although these methods offer rapid ASTs, the universal application of probe molecules to multiple strains and antibiotics remains to be addressed.

Instead of using high-resolution imaging or complicated readout systems, laser speckles may provide an optimal tool for rapid measurement. Laser speckles are a characteristic granular pattern formed by the coherent diffraction of diffusive objects^25,26^. Speckles are often considered as a noise in a coherent imaging system that degrade image contrast^27,28^. However, due to their high sensitivity to the changes in optical phase, speckles have also been exploited to develop sensors for various purposes such as pressure^29^, temperature^30^, and mechanical vibrations^31^. In biomedical applications, laser speckle contrast imaging has widely been used for blood flow measurement^32,33^, microorganism detection^34,35^, and studying drug action on parasites^36^.

Here, we present a rapid AST method based on laser speckle formation (LSF). The irregularity induced by bacteria on agar surface allows the agar plate to function as an optical diffuser. Consequently, the incoming laser beam is diffused resulting in LSF. Because LSF depends on the number of bacteria on the agar plate, speckles develop as the bacteria grow on the agar surface. Therefore, bacterial growth can be detected by quantitatively evaluating the speckles formed. We quantitatively evaluated LSF by calculating the spatial autocorrelation of speckle images and determined bacterial growth by tracking the change in autocorrelation value over time. We performed AST using the proposed method with several combinations of bacterial species and antimicrobial agents. We also validated the proposed method for low bacterial concentrations to demonstrate its sensitivity. The results showed that the proposed method enables rapid AST in 2-4.5 hours.

## Results

### Principle

A conventional method of detecting bacterial growth on agar plates relies on human vision (Fig. 1a). Direct counting of bacterial colonies gives the number of resistant bacteria to the given antibiotic condition. This method requires a long incubation time for the visible growth of each colony, typically 16-20 hours in standard protocols^37^. To circumvent this long lead time, we exploit the phenomenon that densely grown bacterial colonies serve as optical diffusers (Fig. 1b). When an agar plate is illuminated with a laser beam, the colonies deform the wavefront. As light propagates, this results in a speckle pattern. Speckle formation is evaluated quantitatively using angle-averaged spatial autocorrelation (see **Quantitative analysis of LSF** for details). The temporal change in autocorrelation value is used to determine the growth of bacteria at an earlier time point than the detection time based on human vision. The schematic of the setup for the proposed method is shown in Fig. 1c. The setup comprises spatial-filtered laser light, sample stage, and camera (see Methods for details and Supplementary Fig.1 for setup picture).

**Fig. 1.**
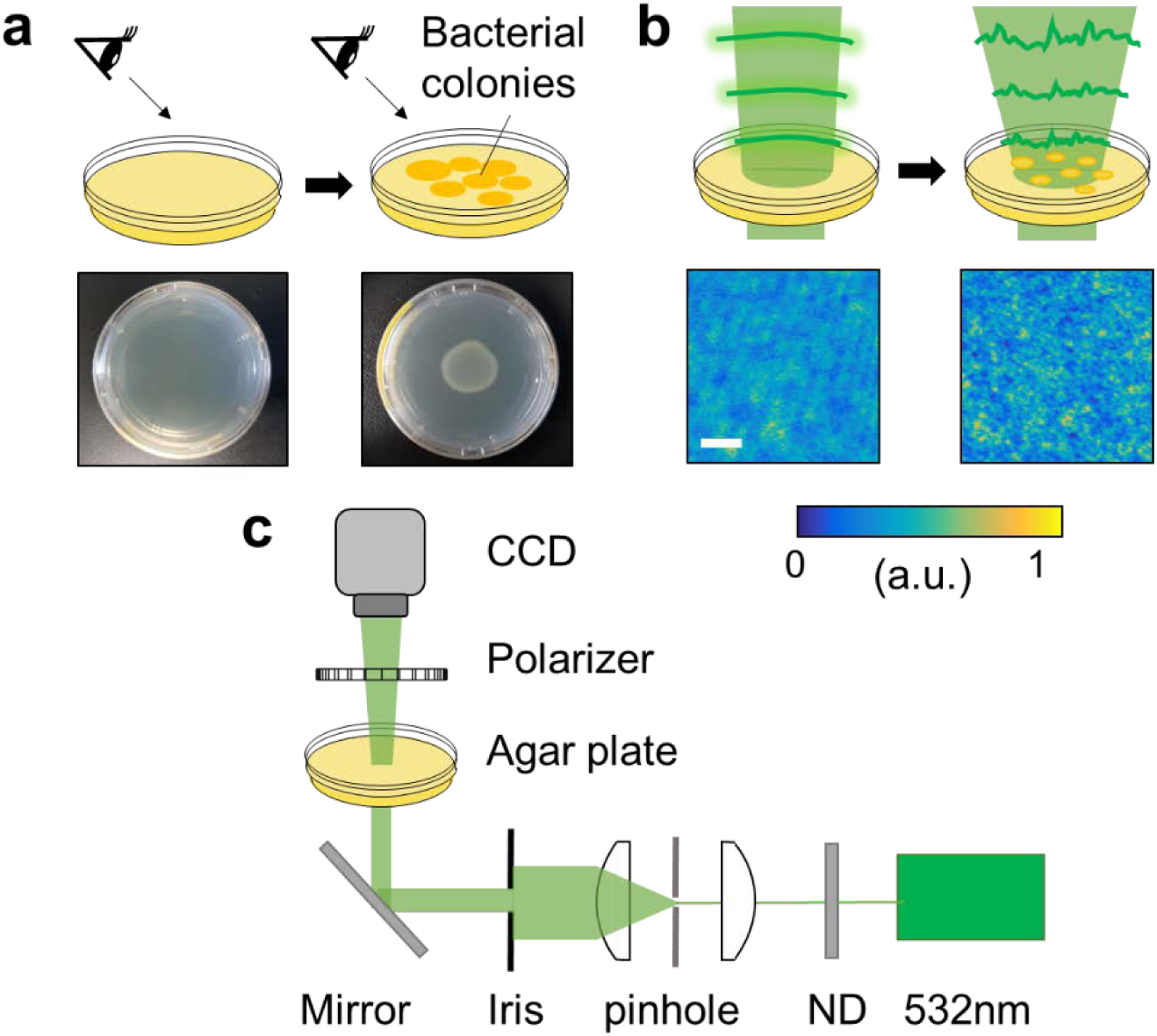
Principle of bacterial colony detection using laser speckle formation (LSF). **a** In conventional methods, bacterial colonies are identified via human vision. It requires a relatively long incubation time to make the colonies visible, typically 16-20 hours in standard methods. **b** Bacterial colonies diffuse incoming light generating speckles. As the colonies grow bigger, the speckles are formed, allowing detection of bacterial growth on agar plates. The scale bar represents 1 mm. **c** Optical setup of the proposed method. Spatial-filtered laser beam passes through the agar plate and the generated speckles are recorded by the CCD camera.

### Bacterial growth and speckle development

The optical sensitivities of human vision, a phase-contrast microscope (CKX53, Olympus, Japan) and speckles were compared using a standard clinical bacterial strain *Escherichia coli* (ATCC 25922). Figure 2a shows the visual development of bacterial colonies. *E. coli* colonies are not observable until two hours and are barely observable after four hours. The colonies are visible after twenty hours, which is the requirement for standard ASTs. In the microscopy images, bacterial growth is more obvious at four hours (Fig. 2b). Each cell forms micro-colonies with increased size from the initial single bacteria. However, the difference in microscope images between zero and two hours is less dramatic since the increase in colony size does not exceed the statistical variation of the initial colony size. The difference between zero and two hours is explicitly visible in speckle images (Fig. 2c). The coarse pattern turns into a fine pattern during the first two hours. The image eventually becomes a clear speckle image with a high contrast at eight and twenty hours. These results show that monitoring bacterial growth using speckle images provides a much faster examination than the conventional method based on human sight. Furthermore, speckle images intrinsically provide a comprehensive approach relative to microscopic images of individual cells.

**Fig. 2.**
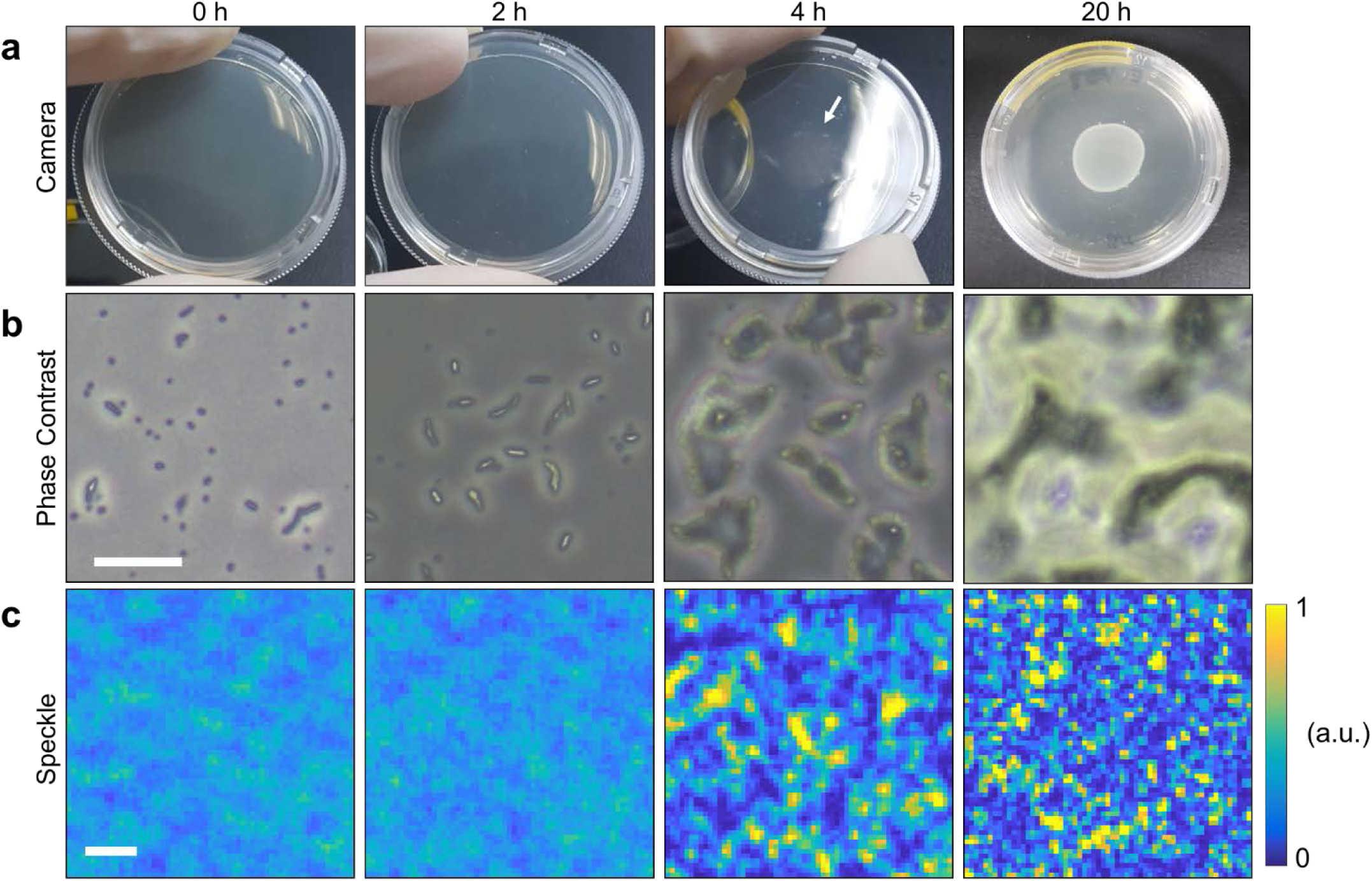
Bacterial images using a camera (human vision), a phase contrast microscope, and speckles. 5 mL of *E. coli* solution of 10^7^ cfu/mL is loaded on the agar plate. **a** Camera pictures of the agar plate at each time. The bacterial colonies are not visible at 0, 2 h and barely observable at 4 h (white arrow). **b** Microscopic view of bacteria on agar plates using a phase contrast microscope. The bacterial colonies are apparently developed at 4 h, while not being clearly recognizable at 2 h. The scale bar represents 50 μm. **c** Speckle images at each time. Speckles start to develop at 2 h. The scale bar represents 50 μm.

### Quantitative analysis of LSF

To evaluate speckle formation by bacterial growth, we used spatial autocorrelation of speckle images^38^. Figure 3a shows the representative camera images for an *E. coli* sample at t_0_ = 0.3 h, t_1_ = 4 h, and t_2_ = 8 h, revealing the development of speckle over time. For each image, the autocorrelation was calculated using the following definition:

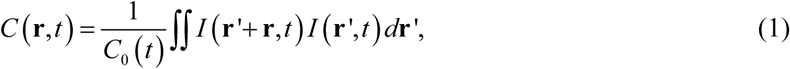

where **r, r’** are position vectors and *C*_0_ (*t*) = ∫ *I*^2^ (**r**’, *t*)*d***r**’. By definition, the autocorrelation for a constant-intensity image is one, and the autocorrelation value decreases when the image intensity differs pixel by pixel. It is clearly seen that the autocorrelation values in Fig. 3b decrease as the speckle develops. Note that the autocorrelation of speckle images does not exhibit a directional tendency. This is because the illumination beam is perpendicular to the sample and camera plane. With this property, the spatial autocorrelation is simplified using the angular average as follows:

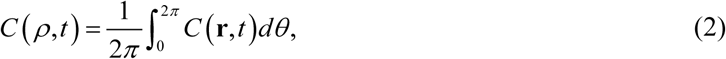

where **r** is represented as polar coordinates: **r** = (*ρ, θ*). The corresponding graph is shown in Fig. 3c.

**Fig. 3.**
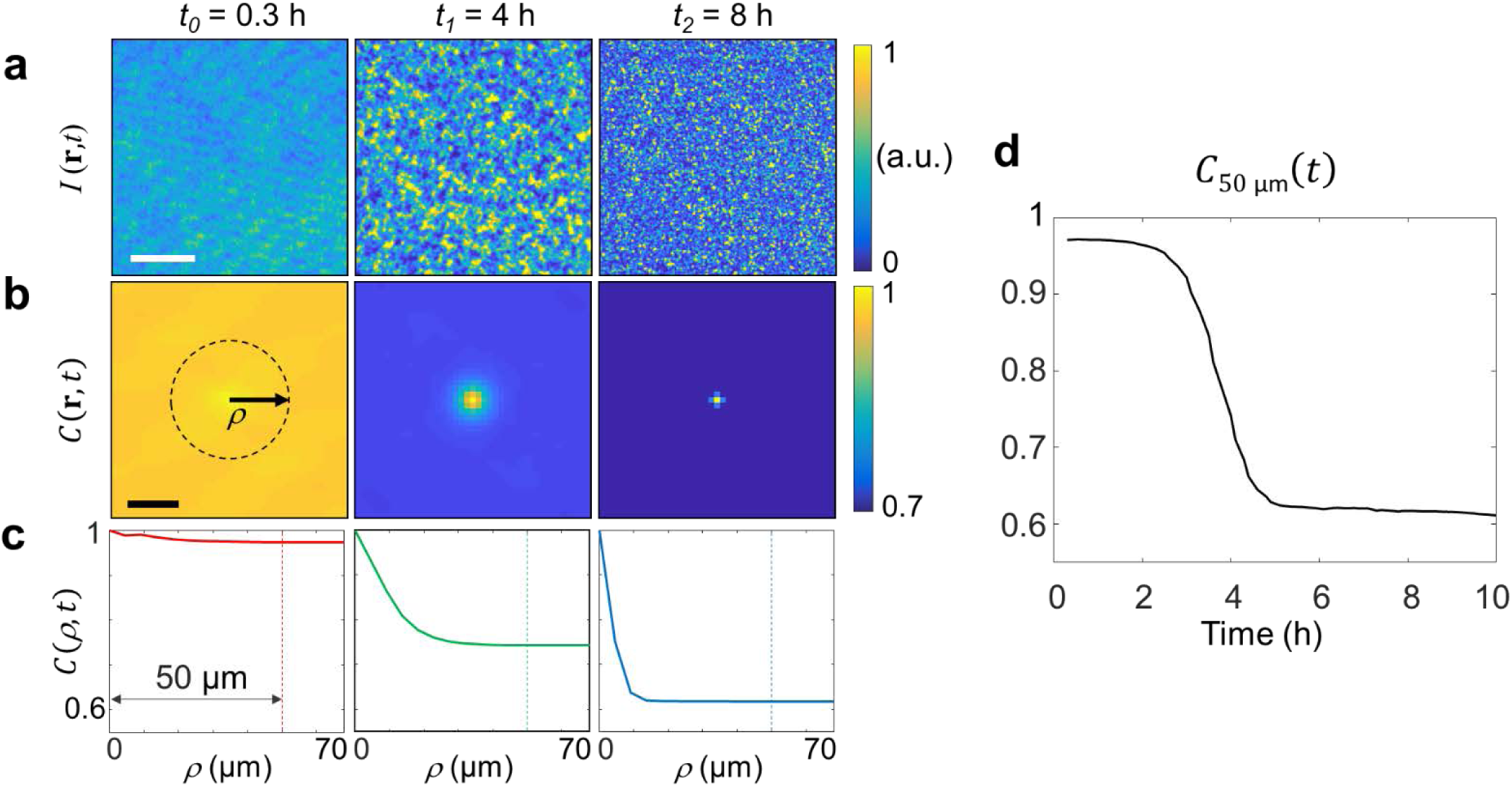
Quantitative evaluation of speckle formation. **a** Speckle images at each time. Speckles are formed as time passes. The scale bar represents 200 μm. **b** The corresponding autocorrelation of each image in a. The autocorrelation not exhibit directional tendency. **c** Angle-averaged autocorrelation graphs for each time. The *C*(*ρ,t*) graph decreases as *ρ* increases, but remains constant after a certain *ρ* value of ten to twenty micrometers. The history of *C*(*ρ,t*) graph is evaluated by tracking its value at *ρ* = 50 μm (dotted lines and arrows), where 50 μm is chosen to be larger than decreasing range. The scale bar represents 40 μm. **d** The graph of angularly averaged autocorrelation value at *ρ* = 50 μm (*C*_50 μm_ (*t*)) over time. The *C*_50 μm_ (*t*) graph has an inverted S-shaped curve, representing the temporal evolution of speckles.

The autocorrelation of speckle images during the entire incubation could be summarised into a single graph with the analysis described below. In Fig. 3c, the decrease of *C* (*ρ, t*) mostly occurred within 10-20 μm of the *ρ* value, and the graph remained almost constant thereafter. This is because the “local property” of image patterns is only valid within the range; in other words, spatial separation of greater than ten to twenty micrometres sufficiently erases the dependence of the autocorrelation value on shift distance. This property allowed us to evaluate LSF using a single parameter, *C* (*ρ*_1_, *t*), with any value of *ρ*_1_ greater than the range. We used the value of *ρ*_1_ = 50 μm that is sufficiently larger than the range (dotted line in Fig. 3c). Following the value of *C* (*ρ*_1_, *t*) over time (denoted as *C*_50 μm_ (*t*)), the time-lapse graph of *C*_50 μm_ (*t*) reveals the history of speckle development in the sample (Fig. 3d). The *C*_50 μm_ (*t*) graph exhibited an inverted S-shape that starts near one and decreases as time passes. The decrease indicated the formation of speckle or equivalently, bacterial growth. The S-shape in the graph matched the well-known curve of bacterial growth^39^, suggesting that the graph represents bacterial growth efficiently. Note that the first time point of the *C*_50 μm_ (*t*) graph was 0.3 h because the time for drying the agar plate and the stabilization of the setup were excluded (see Methods). In addition, the autocorrelation value at *t*_0_ was not exactly one because the output beam was not a perfect plane wave due to the irregularity of the agar plate. Therefore, in the next sections, we used *C*_50 μm_ (*t*) − *C*_50 μm_ (*t*_0_) graph to compensate for the decrease caused by the agar plate, to enable comparison between different agar plates.

### Rapid AST using the proposed technique

AST was performed using the proposed technique in order to demonstrate that the method achieves rapid AST. Three standard clinical strains were used, of which two were gram-negative strains, *Escherichia coli* (ATCC 25922), *Staphylococcus aureus* (ATCC 29213), and one was a gram-positive strain, *Pseudomonas aeruginosa* (ATCC 9027). For antibiotic agents, we used ampicillin for *E. coli* and norfloxacin for the other two strains. Ampicillin is a beta-lactam antibiotic that has excellent activity against *E. coli*^40^. The reference MIC range of ampicillin to *E. coli* is 2-8 μg/ml^41^, and in our experiment, the measured MIC through agar dilution was 2 μg/ml. Norfloxacin is a fluoroquinolone antimicrobial agent that is effective for treating urinary tract infections including infections caused by *P. aeruginosa*^42^, and is also effective against *S. aureus*^43^. The reference MIC range of norfloxacin to *S. aureus* is 0.5-2 μg/ml^41^ and that to *P. aeruginosa* is 0.5-8 μg/ml^44^, where measured MICs were 2 and 8 μg/ml, respectively. For these three combinations of bacterial species and antimicrobial agents, the proposed technique was tested for AST performance. For each combination, two concentrations of antibiotics that are lower and higher than the MIC were tested, and each condition was repeated for ten different samples.

Figure 4a shows the angle-averaged autocorrelation graphs of AST results. Here we used the *C*_50 μm_ (*t*) − *C*_50 μm_ (*t*_0_) graph instead of *C*_50 μm_ (*t*) graph because of the reason discussed at the end of the previous section (see Supplementary Figure 2 for *C*_50 μm_ (*t*) graphs). Graphs of no antibiotics and low antibiotic concentration exhibited inverted S-shapes, whereas those for high antibiotic concentration remain almost constant similar to the control (PBS only). The autocorrelation values between growing and non-growing samples in Fig. 4a are clearly separated in a few hours, demonstrating that the proposed method allows rapid AST.

**Fig. 4.**
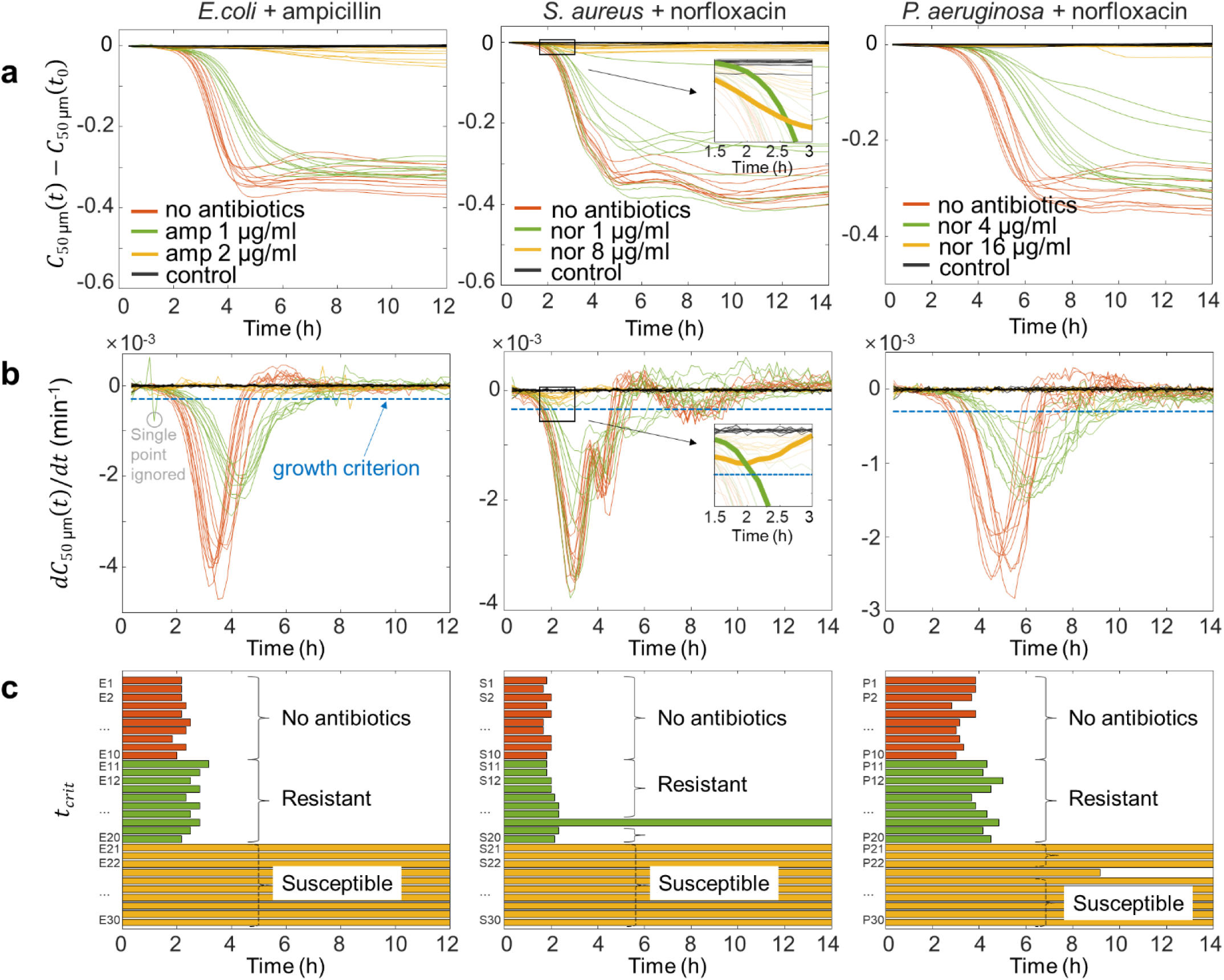
AST results using the proposed method. Three combinations of bacterial species (*E.coli* ATCC 25922, *S. aureus* ATCC 29213, and *P. aeruginosa* ATCC 9027) and antibiotic agents (ampicillin and norfloxacin) are investigated. For each combination, three different concentrations of antibiotics (zero, low and high) are tested beside control (PBS only). The antibiotic concentrations are chosen to be higher and lower than the minimum inhibitory concentration (MIC), where MIC is formerly measured by agar dilution. Each condition is repeated for ten different samples. **a** *C*_50 μm_ (*t*) − *C*_50 μm_ (*t*_0_) graph of each sample. Only samples with antibiotic concentration under MIC have explicit decrease. **b** *dC*_50 μm_ (*t*)/*dt* graph of each sample. The inset indicates that derivative provides a more efficient separation between growing and non-growing samples. The threshold of value -3.5×10^−4^ min^-1^ is used as the growth criterion (blue dotted line). Single outlying points are ignored for criterion (gray circle). **c** Using the threshold, the first time point when graph exceeds the threshold for two consecutive points (*t*_crit_) is shown for every sample. Resistant and susceptible samples are properly identified for most cases.

To quantitatively analyse the AST time, we suggest a growth criterion using a derivative of the autocorrelation graph (Fig. 4b). The reason we used a derivative as the criterion instead of the autocorrelation itself is revealed in the insets of Figs. 4a and 4b for *S. aureus* + norfloxacin. In the inset of Fig. 4a, the green and yellow curves overlap until 2.5 h. The green curve steepens while the yellow curve becomes horizontal during the overlapping range, and the difference in the slope separates the two cases at a much earlier time of 2 h, as can be seen in the inset of Fig. 4b. Therefore, using the derivative as the criterion facilitates a more rapid determination of bacterial growth. We used the threshold with a value of -3.5×10^−4^ min^-1^ as the criterion for bacterial growth (blue dotted line in Fig. 4b). Single outlying points, such as the grey circle in Fig. 4b, were considered as noise and ignored for the growth criterion. Using the threshold value, the first time point when the graph exceeds the threshold for two consecutive points (*t*_crit_) is shown in Fig. 4c for every sample. The resistant and susceptible cases are well separated by *t*_crit_. However, one sample in *S. aureus* + norfloxacin 1 μg/ml was not classified as resistant. We found that the corresponding sample did not develop colonies, which means the sample was susceptible to the given antibiotic condition. Excluding this case, the values of *t*_crit_ of resistant samples were 2.78 ± 0.06 h, 2.11 ± 0.04 h, and 4.52 ± 0.18 h for *E. coli, S. aureus*, and *P. aeruginosa*, respectively. Determining the antibiotic susceptibility in 2-4.5 hours, the results show that the proposed technique reduces the time needed for AST by five to ten times compared to the 16-20 hours for the standard protocol of agar dilution.

### Sensitivity for low-density samples

We show that the proposed method is also applicable for samples with low-density or individual colonies. The ability to detect a few colonies is essential because only a portion of bacteria may survive at some moderate concentration of antibiotics. To demonstrate the low-density sensitivity, the proposed technique was tested at different bacterial concentrations: 10^7^, 10^6^, 10^5^, and 10^4^ cfu/ml. Figure 5a presents the camera image of each concentration at eight hours after the sample was loaded. The samples with high concentrations, 10^7^ and 10^6^ cfu/ml, showed well-developed speckle images. In contrast, the images of low concentration samples, 10^5^ and 10^4^ cfu/ml, exhibit individual colony spots rather than speckles. Figure 5b shows the results of samples with different bacterial concentrations using the proposed method. While all bacterial samples have an inverse peak in the *dC*_50 *μ* m_ (*t*)/*dt* graph, the peak size decreases when the bacterial concentration is low. Using the aforementioned criterion, the proposed technique detects bacterial growth at concentrations as low as 10^5^ cfu/ml. As the standard protocol of AST requires 10^7^ cfu/ml as the initial concentration, this result indicates that 1% of the surviving bacteria can be detected within the standard AST sample.

**Fig. 5.**
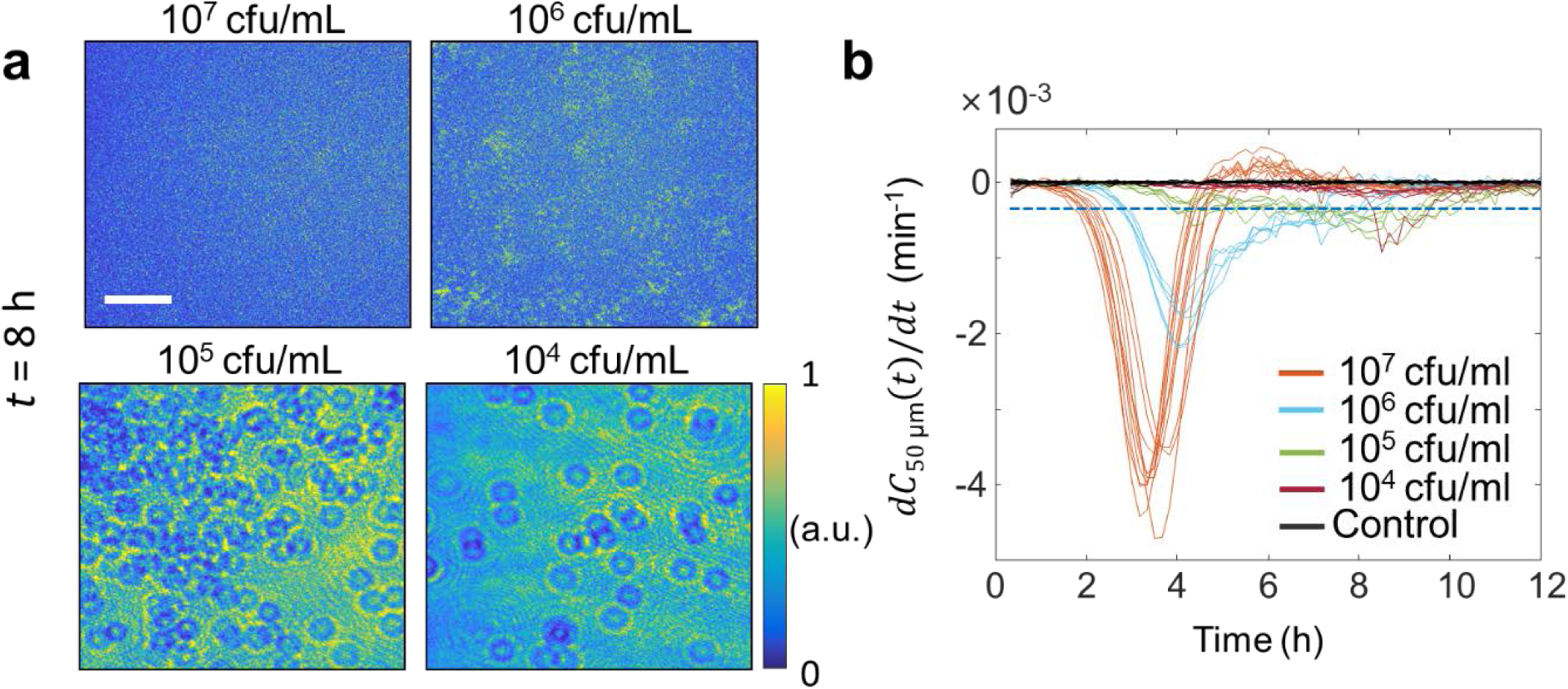
The proposed method is demonstrated under four different concentrations of *E. coli* with no antibiotics. **a** Camera images for each bacterial concentration at *t* = 8 h. Speckle are well developed with samples of 10^7^, 10^6^ cfu/ml. Lower concentrations of 10^5^, 10^4^ cfu/ml exhibit individual colonies on the images rather than speckles. The scale bar represents 1 mm. **b** The *dC*_50 μm_ (*t*)/*dt* graph for different bacterial concentrations. For all tested concentrations, different concentrations are distinguishable, while samples of same concentration have similar graphs. Samples of lower concentration have a smaller inverse peak in the *dC*_50 μm_ (*t*)/*dt* graph. Using the criterion mentioned before, the method could detect the growth of bacteria with concentration as low as 10^5^ cfu/ml.

## Discussion

The biggest benefit of the proposed technique is that it shares the sample preparation protocols with the agar dilution method, which is the standard method for solid-based AST^45^. The proposed method involves inoculation of bacteria on antibiotic-containing agar plates to determine susceptibility, which is identical to that of agar dilution^2^. Since subtle differences in testing methods can influence AST results^45^, the preservation of protocol in the present method that circumvents this problem is a great advantage for point of care applications. Therefore, the current method is readily available for institutions, hospitals or laboratories that use agar dilution for various purposes.

The proposed method determines bacterial susceptibility in 2-4.5 h, which is significantly faster than the standard AST methods (∼16-20 hours) ^7,37^ and commercialised automated systems (∼8 h)^7-10^. There are several factors contributing to this rapidity. First, speckles are sensitive to both amplitude and phase changes induced by bacteria. Since bacterial colonies have been reported to exert phase modulation^46^, speckle patterns can sensitively respond to changes of optical path length in bacterial samples better than amplitude-dependent detection such as camera-vision based AST (∼ 3.5 h)^47^. Second, the proposed technique averages the response over a large population, unaffected by individual variation of microorganisms. At certain antibiotic concentrations, some percentage of bacteria may be resistant whereas the rest is susceptible to the antibiotic. MIC_50_, for example, is the antibiotic concentration that kills 50 percent of the isolate of a test species^48^. In this situation, microscopy-based single-cell analysis may report both resistant and susceptible depending on the choice of target bacteria, which forces investigation of multiple spots. In contrast, the proposed technique determines susceptibility based on the information across the entire isolate on the agar plate, compensating for variations depending on specific colonies. This enables more efficient measurement of bacterial responses, which supports that the proposed technique has an equal or shorter AST time compared to a previous direct single-cell imaging method (∼4 h)^12^. Finally, the current method performs much faster AST than the method described in our previous paper^49^. The previous paper also used speckle correlation but analysed the temporal change in images instead of spatial correlation. The main obstacle of using temporal change was that bacterial movement overlapped with the dehydration of agar gel. The current method can deviate such issues because dehydration does not affect spatial autocorrelation.

Although the proposed method is simple and fast, several requirements should be satisfied to perform the present method. The coherence length of the laser beam should be sufficient to produce a speckle contrast and the input laser beam should be set as a plane wave. Mechanical movement during camera exposure should be minimised for accurate measurement of LSF. These requirements should be fulfilled to guarantee the clarity of speckle images.

The current technique has a different purpose and capability from previous speckle-based microorganism detections. In contrast to dynamic speckle analysis^26,50-52^, which focuses on the temporal differences in images of speckle patterns, the current technique quantifies the autocorrelation of speckles and follows the history of this value. One advantage of the method proposed here is that it is not affected by mechanical movements between frames because an image shift does not interfere with spatial autocorrelation. Another speckle-based microorganism detection method involves analysis of laser-induced Fresnel^53^ or speckle scatter patterns^34^ in bacterial colonies. This method focuses on scatter patterns by a single colony to identify the bacterial species, in contrast to the proposed method that examines speckles of high-density colonies to determine cell survival at given antibiotic conditions.

In this work, we demonstrated the eligibility of the proposed method for rapid AST by testing several combinations of bacterial species and antimicrobial agents. The present method, in principle, operates well as long as the standard agar dilution test is applicable because speckle formation is not dependent on the bacterial species or antibiotics. The proposed method using LSF has many benefits in multiplexing and parallelizing due to its simple, robust, and low-cost configuration that requires much fewer components compared to high-resolution lens systems. A high-throughput automated system would be helpful for its further study and broad application. We anticipate that the proposed method using LSF could bring an innovative advancement in rapid AST tools.

## Methods

### Bacterial species and culture

We used bacterial strains *Escherichia coli* (*E. coli*, ATCC 25922, Korean Collection for Type Cultures (KCTC), Republic of Korea), *Staphylococcus aureus* (*S. aureus*, ATCC 29213, Korean Collection for Type Cultures (KCTC), Republic of Korea), and *Pseudomonas aeruginosa* (*P. aeruginosa*, ATCC 9027, Korean Collection for Type Cultures, Republic of Korea). All strains were purchased from the corresponding suppliers. The culture media for each species were adopted as listed on the species specification; *E. coli* and *P. aeruginosa* were cultured in Nutrient broth (ingredients purchased from BD Biosciences, USA), and *S. aureus* was cultured in Luria-Bertani broth (ingredients purchased from BD Biosciences, USA). The manufacturing procedure for culture media strictly followed the protocol from KCTC.

The bacterial species were cultured overnight and were then diluted to the desired concentration for the experiment. The bacteria were centrifuged for 10 minutes at 2000 rpm. The medium was then aspirated and the bacteria collected at the bottom were suspended in phosphate buffered saline (PBS, 1X, 0067M PO4, Hyclone Inc., USA). The bacterial concentrations were adjusted by measuring optical density (OD) with a spectrophotometer (UV-1280, Shimadzu Inc., Japan) at λ = 600 nm. The viable count of the adjusted bacterial solution was confirmed through dilution and incubation on agar plates^37^. Bacterial solutions were initially adjusted to 0.15-0.2 OD (1×10^8^ CFU/ml), followed by ten-fold dilutions until the desired concentration was obtained.

### Agar plate and sample preparations

The procedure for making agar plates followed the protocol from KCTC. Appropriate culture broth components and agar powder (7.5 g for total volume 500 ml) were mixed with distilled water and autoclaved. The solution at 70°C was distributed into Petri dishes (40 mm in diameter, TPP 93040, Sigma-Aldrich, Germany) (3 ml per dish). The agar plates were cooled and dried on a clean bench for thirty minutes with the fan off. For antimicrobial susceptibility tests, ampicillin (A0166, Sigma-Aldrich Inc., USA) and norfloxacin (N9890, Sigma-Aldrich Inc., USA) were chosen as the test antimicrobial agents. The antimicrobial stock solution of concentration 100 mg/ml was prepared by diluting with phosphate buffered saline and was stored in a refrigerator at 4°C. To prepare the antibiotic agar plates, the desired amount of the stock solution was added to the autoclaved agar solution at 50°C. The agar solution with antibiotics was solidified on a fan-off clean bench for thirty minutes with the cover ajar.

The manufactured agar plates were stored in a refrigerator at 4°C. For sample preparation, agar plates were taken out of the refrigerator just before use. The bacterial solution was loaded on the agar plate, and evenly spread at the centre with a diameter of 2 cm. As control, PBS solution was used instead of a bacterial solution. After drying on a clean bench (fan operating) for about ten minutes, the agar plate was placed upside down on the sample stage with a temperature controller (35°C). Excluding ten minutes for drying and another ten minutes for plating, the measurement started twenty minutes after the bacterial solution was loaded.

### Optical setup

The optical system was designed with a spatially filtered laser beam, a neutral density filter (denoted as ND in Fig. 1c), an iris, a polarizer, and a camera. A laser beam (Cobolt Samba 532 nm laser, 25 mW, Cobolt, Inc., Sweden) was spatially filtered with a pinhole (15 μm) and its intensity was adjusted using ND. The diameter of the beam was set to be 15 mm using the iris. The agar sample was placed in a custom-built aluminium moulding with a temperature controller. The beam illuminated the centre of the agar plate where bacteria were seeded. After the sample stage, a linear polarizer (LPVIS100, Thorlabs Inc., USA) was added for better visibility. Finally, a CCD camera (Lt365R, Lumenera Inc., USA) was mounted vertically facing down toward the sample with the distance from sample being approximately 7 cm. The size of a pixel in the camera was 4.54×4.54 μm^2^, and the whole image was 1024×1024 pixels. For the main vertical structure, 1.5-inch damped posts (DP14A, Thorlabs Inc., USA) were used to achieve mechanical stability.

